# Aperiodic dynamics track cortical state shifts during sleep K-complexes

**DOI:** 10.64898/2026.09.23.753632

**Authors:** Elizabeth S Kaplan, Jonathan Schindler, Bradley Voytek

## Abstract

Sleep microarchitecture is defined by brief electrophysiological events such as K-complexes (KCs), which are traditionally interpreted as discrete oscillatory phenomena involved in sleep maintenance, sensory gating, and cortical down-states. However, electroencephalography (EEG) also contains aperiodic, broadband activity that reflects non-oscillatory population dynamics and varies systematically across sleep stages. Because KCs are large, low-frequency transient waveforms, they also manifest across a broader frequency range, which creates a bidirectional measurement problem in which discrete events and aperiodic activity are difficult to disentangle. This is because aperiodic dynamics alone can generate fluctuations that resemble KCs, while conversely, the KC waveform itself can inflate the low-frequency end of the spectrum, steepening the fitted exponent even when the underlying aperiodic state is unchanged. These confounds blur what constitutes a real KC and a real aperiodic change in the presence of a KC. A KC-locked change in the aperiodic exponent could therefore reflect a genuine cortical state shift, a measurement artifact from the waveform biasing the spectral fit, or both. Here, we use time-resolved spectral parameterization of overnight sleep EEG to characterize aperiodic dynamics around expert-annotated KCs, with a series of controls designed to separate genuine aperiodic state changes from waveform artifacts. A component of the KC-locked aperiodic change survives these controls, suggesting that KCs are accompanied by a genuine shift toward a higher-exponent, inhibition-dominated cortical state. These findings establish aperiodic activity as a meaningful index of state dynamics during KCs and highlight measurement confounds that any event-locked spectral analysis of large transients must consider.

**Significance Statement:** The aperiodic component of neural activity is an increasingly used marker of brain state, yet accurately measuring it around transient events is difficult. Sharp waveforms create broadband power and can masquerade as changes in the underlying dynamics. We address this problem using the sleep K-complex, the largest event in healthy human EEG. Combining four complementary strategies, we disentangle waveform-induced spectral distortion from genuine modulation of aperiodic activity, and show for the first time that K-complexes are accompanied by a transient shift in the aperiodic exponent. Beyond this finding, our framework provides a general method to control for biases that waveform shape introduces into spectral estimates, informing an active debate over how transient events are detected in neural signals.

## 1 INTRODUCTION

Sleep is essential for memory, mood, and neural plasticity (Miletínová and Bušková, 2021; Sulaman et al., 2024). Coordinated neural activity supporting sleep is captured with electroencephalography (EEG) (Klinzing et al., 2019). Traditionally, sleep EEG is interpreted through oscillations within canonical frequency bands, linked to distinct physiological and cognitive processes (Tononi and Cirelli, 2006; Cohen, 2017; Cole and Voytek, 2017) that vary systematically with sleep stages (Passaro and Poltorak, 2024). Within NREM Stage 2 (N2), a prominent event is the K-complex (KC), a large biphasic waveform: a brief positive deflection, a high-amplitude negative component, then a later positive phase (Colrain, 2005). Expert annotation remains the gold standard, yet the defining criteria are ambiguous. The American Academy of Sleep Medicine defines a KC as a “well-delineated negative sharp wave immediately followed by a positive component standing out from the background EEG” (Berry et al., 2023). Scorer reliability is correspondingly limited (Lajnef et al., 2017), as KC morphology varies across individuals, states, and clinical populations (Gandhi and Emmady, 2023), leaving near-identical events inconsistently labeled (Figure 1A, B).

**FIGURE 1.**
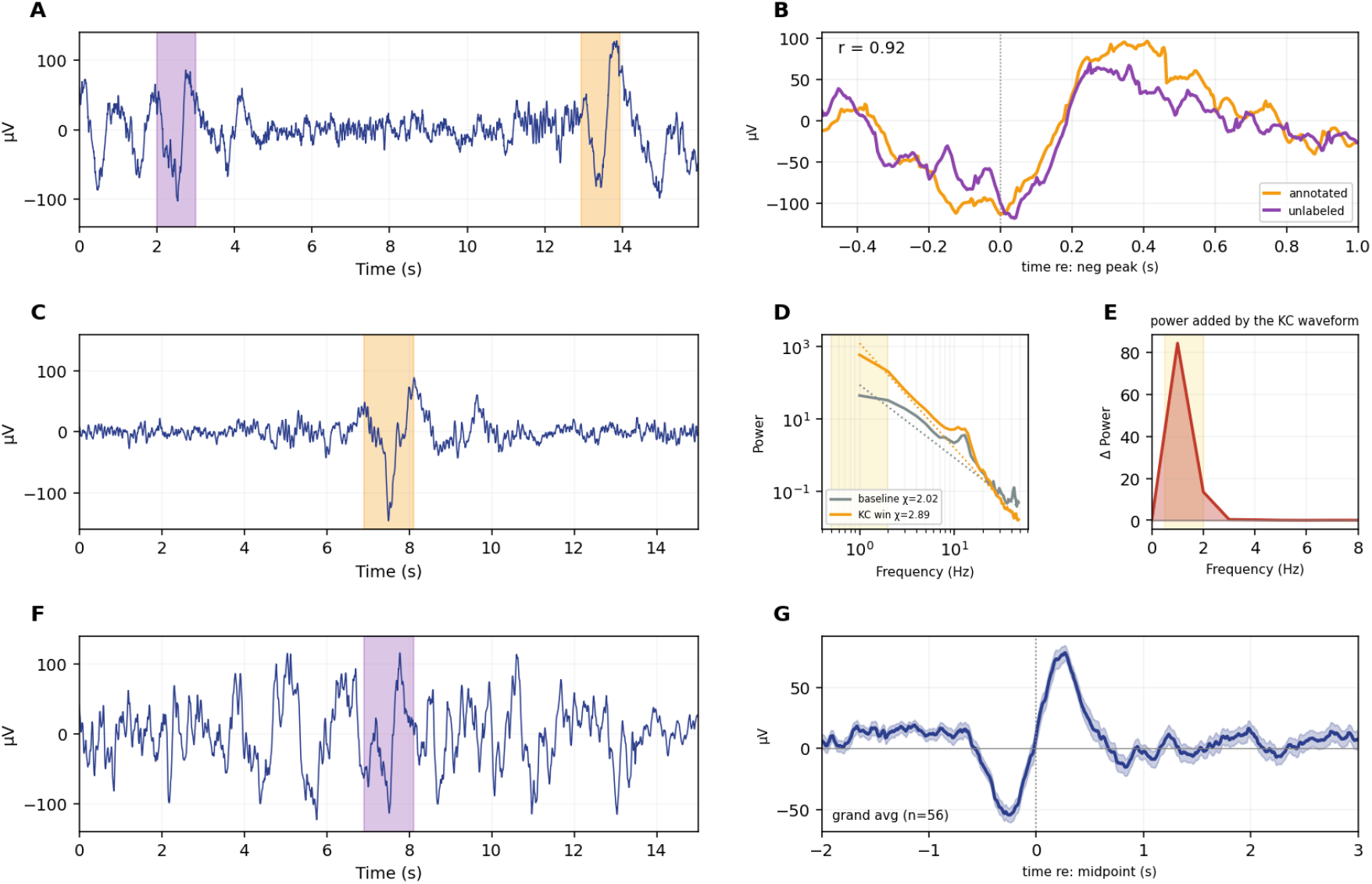
Ambiguity between discrete KCs and aperiodic activity. (A) Representative C3 trace from the MASS dataset, containing two morphologically similar waveforms: one expert annotated as a KC (highlighted, orange), one unlabeled (highlighted, purple). (B) Annotated (orange) versus unlabeled (purple) event overlaid, time-locked to the negative peak. The two events are highly correlated (r=0.92), indicating that the unlabeled event is morphologically identical to the annotated KC. (C) C3 trace from MASS data showing a single annotated KC (highlighted, orange). (D) Power spectrum computed in a baseline window (gray, dotted line: aperiodic fit, χ = 2.02) versus a window centered on the KC (orange, χ = 2.89), showing the exponent steepens when the fit window contains the event. (E) Power added by the KC waveform as a function of frequency, isolating the low-frequency band (∼1-2 Hz; shaded, yellow), where the waveform’s spectral contribution is concentrated. (F) Trace from a 30-minute purely aperiodic simulation, showing a KC-like event (shaded; purple) identified by MT-KCD despite containing no true oscillatory KC in the simulated signal. (G) Grand average waveform of all MT-KCD detections in purely aperiodic signal (n=81), which reproduces the canonical biphasic KC morphology, demonstrating that aperiodic activity alone can generate events that pass KC detection criteria.

This variability may reflect more than imprecise definitions. KCs are typically treated as discrete oscillatory events, but may instead emerge within broader fluctuations in population activity. One candidate source is the aperiodic component of EEG, the broadband, non-rhythmic 1/f structure co-occurring with oscillations (Donoghue et al., 2020). Aperiodic activity reflects physiology including excitation/inhibition (E/I) balance and global changes in neural state (Gao et al., 2017; Preston et al., 2026). The aperiodic exponent changes over the night, tracking sleep stages, transitions, and responses to stimuli (Ameen et al., 2025; Lendner et al., 2024; Rosenblum et al., 2024).

Time-resolved estimates make it possible to examine aperiodic activity around individual KCs, but the two are difficult to disentangle. Physiologically, KCs are linked to cortical downstates and brief inhibitory shifts (Cash et al., 2009). Such E/I shifts manifest as a steepening of the aperiodic exponent (Gao et al., 2017), suggesting KCs may coincide with genuine changes in the aperiodic background. Definitionally, a KC is a large low-frequency transient. When estimated in a window containing a KC, the event’s power inflates low frequencies and biases the fit even when the underlying state is unchanged (Figure 1C-E). Because a real state shift and this artifact push the exponent in the same direction, a KC-locked change cannot, on its own, distinguish the two.

This ambiguity is exacerbated because aperiodic activity alone can produce KC-like waveforms. In a purely aperiodic simulation, the Multitaper K-Complex Detector (MT-KCD; Oliveira et al., 2020) detects 81 events whose grand-average reproduces the canonical KC morphology (Figure 1F, G). A parallel observation holds for hippocampal sharp-wave ripples, detectable in purely aperiodic signals (van Schalkwijk and Helfrich, 2026). Subsequent work argues that such surrogates inflate false positives through adaptive thresholds, and that genuine events remain distinguishable by their temporal and spectral structure (Kragel, 2026). Both sides argue from aggregate detection rates rather than individual events, leaving the status of any single event unresolved.

Direct evidence linking KCs to aperiodic activity is limited. Ameen et al. (2025) found auditory-evoked exponent increases that were larger when a KC was elicited but also present without one, suggesting the shift and the waveform are distinct. But that work was locked to external stimuli, not spontaneous KCs, and did not separate a genuine state change from the waveform artifact. This matters because the features that define a KC, its amplitude and low-frequency morphology, are what most bias a spectral fit.

Here, we address this gap. We apply time-resolved spectral parameterization to overnight EEG, characterizing the aperiodic exponent around spontaneous, expert-annotated KCs. We then apply controls designed to separate genuine aperiodic state changes from waveform-driven artifacts. Together, they ask whether part of the KC-locked exponent increase survives correction, and more broadly show how discrete waveforms bias event-locked spectral estimates, a confound any analysis of aperiodic activity around large transient events must consider.

### 2 MATERIALS AND METHODS

*Participants and data*. Whole-night polysomnography (PSG) recordings were obtained from the SS2 subset of the Montreal Archive of Sleep Studies (MASS; O’Reilly et al., 2014), comprising 19 healthy young adults (11 F; age 23.6 ± 3.7 years, range 18–33). Recordings were acquired with 19 EEG electrodes placed according to the international 10–20 system, together with four EOG, one bipolar EMG, and one ECG channel, at a sampling rate of 256 Hz. Expert-scored sleep stages and expert-annotated KCs, scored using a minimum duration of 0.5 s and a minimum peak-to-peak amplitude of 75 µV, were provided with the dataset. KC annotations were scored on the C3 electrode.

### Preprocessing

The C3 signal was extracted, and N2 periods were isolated for each participant using the expert staging annotations. N2 intervals separated by less than 0.5 seconds were concatenated into a continuous N2 recording. To avoid discontinuities at the concatenation boundary, a 0.5 second half-cosine taper was applied on either side of each segment boundary. Preprocessing was performed in MNE-Python (Gramfort et al, 2013). The continuous N2 data for each subject was band-pass filtered from 0.1 to 100 Hz and notch-filtered at 60 Hz using a zero-phase filter. Ocular artifacts were removed using independent component analysis (ICA). Following ICA, the data were re-referenced to the contralateral mastoid.

### Time-resolved spectral parameterization

To characterize the temporal evolution of aperiodic activity across the night, a time-resolved spectral parameterization was applied to each subject’s C3 recording (Cellier, 2026). The signal was divided into overlapping 2 second windows with a 0.25 second step (75% overlap). For each window, the power spectral density (PSD) was estimated using the multitaper method (mne.time_frequency.psd_array_multitaper) with a frequency bandwidth of 4 Hz and parameterized using the specparam toolbox (SpectralGroupModel; Donoghue et al., 2020) in the fixed aperiodic mode, with peak width limits 1–12 Hz, maximum of 5 peaks, minimum peak height 0, and peak threshold of 2 standard deviations, yielding time-resolved estimates of the spectral exponent. Model fit was quantified per window using the goodness-of-fit R^2^; windows with R^2^ < 0.9 were excluded from further analysis. The model was fit over a systematically varied lower frequency bound, with the upper bound (45 Hz) and all other parameters held constant, so that the effect of progressively excluding the KCs low-frequency power could be assessed (Figure 4E). We report statistics across three primary fit ranges: a broadband (1-45 Hz), which includes the KCs spectral footprint, and two highband control ranges (10-45 and 20-45 Hz) that progressively exclude the low-frequency power deposited by the KC waveform. The 20-45 Hz range additionally excludes the sigma/spindle band (11-16 Hz).

### Event-locked aperiodic dynamics

Because KC onsets are not well-defined, and expert annotations from the MASS dataset fall at variable positions relative to the waveform (SD 430 ms; Figure S1), each KC onset was first re-aligned to the negative peak of the C3 signal within the 1 second following the mark. This amplitude-based alignment is deliberately conservative: by locking onto the largest low-frequency deflection it maximizes any waveform-driven bias in the fit, so an exponent change that persists in the high-band controls cannot be attributed to the waveform. The time-resolved exponent trace was epoched from −10 to +10 s around each KC’s negative peak in each fit range, and averaged across events within-then across-subjects to form the grand-mean trace. The event-locked deflection was defined as the peak of this grand-mean trace relative to the far-field N2 baseline (mean exponent at |t| > 3 s). For the amplitude-coupling control, a per-event deflection was computed for each KC individually as the exponent in the 1 s window at the negative peak minus the mean exponent of two baseline windows at ±5 s, and these per-event deflections were averaged within and then across subjects.

### Isolating genuine aperiodic change from waveform artifact

The KC-locked increase in the exponent could reflect either a genuine change in the aperiodic background or contamination from the waveform itself, which deposits power at the low-frequency end of the spectrum and can artificially steepen the fitted exponent. To distinguish these possibilities, we applied three complementary controls, each targeting the confound using a different approach. Together, they test how much of the observed change in the exponent around the event is attributable to the waveform artifact versus a genuine shift in the underlying aperiodic exponent, and whether the artifactual contribution scales with the waveform’s amplitude.

First, we refit the aperiodic model above the KC’s frequency footprint. Because a KC deposits its power at low frequencies, the model was refit over two highband ranges (10–45 and 20-45 Hz), where the KC contributes negligible power; a deflection that persists in these ranges is much less likely to be explained by the waveform and more likely indicates a change in the aperiodic background.

Next, we estimated the exponent change expected from the waveform alone using an injected-waveform artifact floor, which measures the exponent change produced purely by the KC waveform’s spectral signature in the absence of any true change in the aperiodic background. The artifact floor serves as the baseline that real KC-locked exponent changes must exceed to be interpreted as genuine state changes. In order to reduce bias introduced by expert annotations, we first passed each subject’s continuous N2 recording through MT-KCD. Only KC-events that were identified by both experts and the detection algorithm were included in our analyses. An empirical KC template was built by averaging the waveforms of isolated KCs (no other KC or spindle within 10 seconds), aligned to the negative peak and baseline-corrected to a pre-event window.

Isolating the waveform’s contribution requires a background that preserves the real spectral structure of the N2 signal but contains no time-locked aperiodic dynamics, thus this template was injected at regularly spaced positions into a phase-randomized Fourier surrogate of each subject’s own N2 recording (Prichard and Theiler, 1994; Lancaster et al., 2018). Each surrogate was generated by computing the discrete Fourier transform of the continuous N2 signal, replacing the phase at each frequency with a value drawn independently and uniformly from [0, 2π], while leaving magnitudes unchanged, and applying the inverse Fourier transform. This assures that the spectral content of the surrogate data are identical to that of the real N2 for each subject. This approach follows the surrogate-matching logic emphasized for event detection in aperiodic signals (Kragel, 2026). The empirical KC template was scaled to each event’s negative-peak amplitude. The artifact floor was averaged across many independent background segments rather than a single realization. The same time-resolved parameterization and event-locked extraction used for real KCs were then applied to the injected events, yielding an artifact floor trace directly comparable to the real one. The floor defines the value that the real KC-locked change must exceed to indicate a genuine change in the aperiodic background.

Finally, we tested whether the deflection scaled with waveform amplitude. As a waveform-independent check, each KC’s negative-peak amplitude was correlated across events with its exponent deflection, pooled across events and quantified with Pearson’s correlation. A positive relationship (larger KCs producing larger deflections) is the signature of waveform contamination, whereas amplitude-independence is consistent with a genuine state change.

### Waveform Regression

To further quantify how KC waveforms uniquely contribute to the exponent dynamics, we applied a method developed to separate aperiodic changes from spectral distortions introduced by event-related potentials in real data (Gyurkovics et al., 2022). This approach removes the event waveform directly from each analysis window before spectral parameterization. For each KC, the empirical KC template was scaled in amplitude and regressed out of the negative-peak window by ordinary least squares, yielding a residual window from which the waveform had been removed while the surrounding activity was preserved. The aperiodic exponent was then re-estimated on the residual window in each fit range, and its deflection relative to baseline was compared with the deflection of the original, unregressed window. A deflection that persists after the waveform is removed reflects a change in the aperiodic background, whereas one that is eliminated reflects contamination by the waveform.

### Statistical Tests

Subject-level event-locked traces were averaged to a grand mean, and the deflection at the KC negative peak was tested against zero with one-sample t-tests, separately for the broadband (1-45 Hz) and highband (10-45 and 20-45 Hz) fits. To examine the relationship between event amplitude and exponent deflection, the within-subject Pearson’s correlation was computed, and the Fisher z-transformed correlations were tested against zero across subjects. Differences in this coupling between fit ranges were tested with paired t-tests. For the artifact floor, the difference in deflection between real KCs and amplitude-matched template injections into the spectrum-matched surrogate was evaluated with a bootstrap over subjects (10,000 resamples), and 95% confidence intervals were reported for each fit range. A deflection was considered to exceed the floor if the lower bound of the interval exceeded zero. For the waveform regression control, the deflection of the residual, waveform-removed window was tested against zero with one-sample t-tests. Analyses were performed in Python using MNE-Python, NeuroDSP (Cole et al., 2019), SciPy (Virtanen et al., 2020), and the specparam toolbox (Donoghue et al., 2020).

## 3 RESULTS

Time-resolved spectral parameterization around expert-annotated, isolated KCs revealed a sharp, rapid increase in the aperiodic exponent peaking near KC onset (Figure 2B). This increase was present across a ladder of frequency ranges (Figure 2A). In the event-locked trace, it peaked Δχ ≈ +0.53 above the N2 baseline in the broadband (1–45 Hz) fit, and at Δχ ≈ +0.45 and Δχ ≈ +0.33 (10–45 and 20-45 Hz) in the highband fits. The presence of a clear peak in the highband fits, above the KC’s spectral footprint, provided an initial indication that the exponent increase was not solely attributable to spectral leakage introduced by the event’s waveform. The event-locked spectrogram confirms that KC power is concentrated at low frequencies, rising sharply below ∼4 Hz around the negative peak, with little change at higher frequencies (Figure 2C).

**FIGURE 2.**
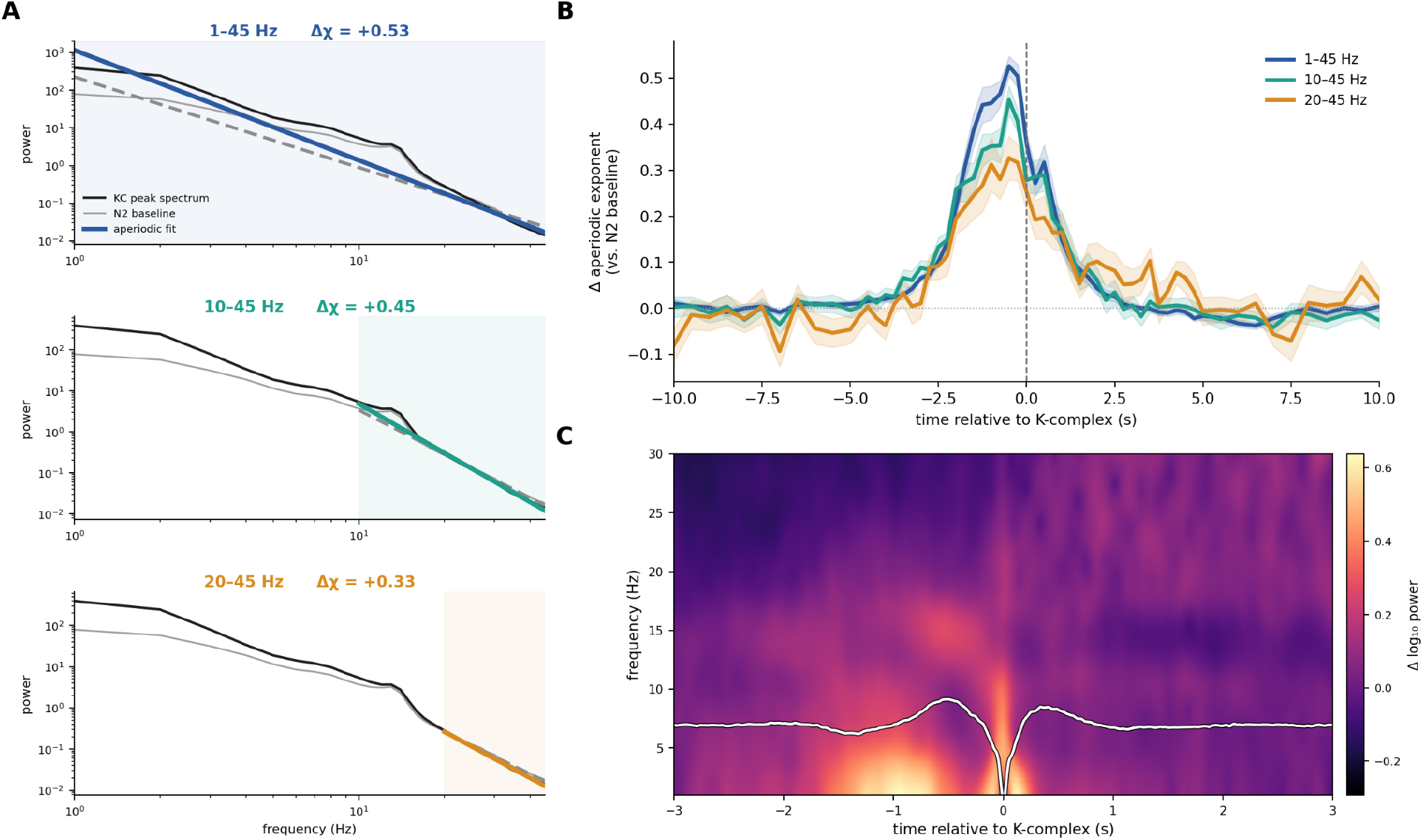
Time-resolved aperiodic exponent dynamics around KCs across fit ranges. (A) Grand-mean power spectra at the time of maximal aperiodic exponent (black) and during N2 baseline (gray dashed), with the aperiodic fit (colored) shown for each of the three fit ranges. Δχ is the peak increase in exponent relative to N2 baseline, matching the trace maxima in (B): +0.53 (1–45 Hz), +0.45 (10–45 Hz), +0.33 (20–45 Hz). (B) KC-locked changes in aperiodic exponent (relative to N2 baseline) as a function of time from KC negative peak, for each fit range. All three show a sharp increase peaking near the KC negative peak, with the effect attenuating as low frequency content is progressively excluded. (C) KC-locked time-frequency spectrogram (Morlet wavelet, n_cycles = f/2) showing that power rises sharply below ∼4 Hz around the KC negative peak (overlaid trace, white), with comparatively little changes at higher frequencies.

Next, to quantify the contribution of the waveform alone, a phase-randomized surrogate was generated from each subject’s N2 recording (Figure 3A) in order to preserve the spectral distribution (Figure 3B). Next, the empirical KC template (Figure 3C) was injected into the surrogate, and the resulting combined signal, referred to as the surrogate floor, was passed through an identical pipeline of time-resolved, event-locked spectral parameterization. In the broadband fit, the real KC-locked deflection (peak Δχ ≈ 0.53) exceeded the surrogate floor (Δ ≈ 0.18), leaving a residual of Δχ ≈ 0.35 not attributable to the waveform (Figure 3D; bootstrap 95% CI [0.29, 0.40], p<0.001). The same relationship was observed in both highband fits. The 10-45 Hz deflection (≈0.45) exceeded the floor (≈ 0.18; residual Δχ ≈ 0.27, 95% CI [0.18, 0.36], p < 0.001; Figure 3E), as did the 20-45 Hz deflection (≈ 0.33 vs 0.08; residual Δχ ≈ 0.25, 95% CI [0.16, 0.34], p = <0.001; Figure 3F). Because the phase-randomized surrogate is stationary in its aperiodic component by construction, the portion of the deflection exceeding the floor reflects a genuine change in the aperiodic background rather than the KC waveform contaminating its own measurement.

**FIGURE 3.**
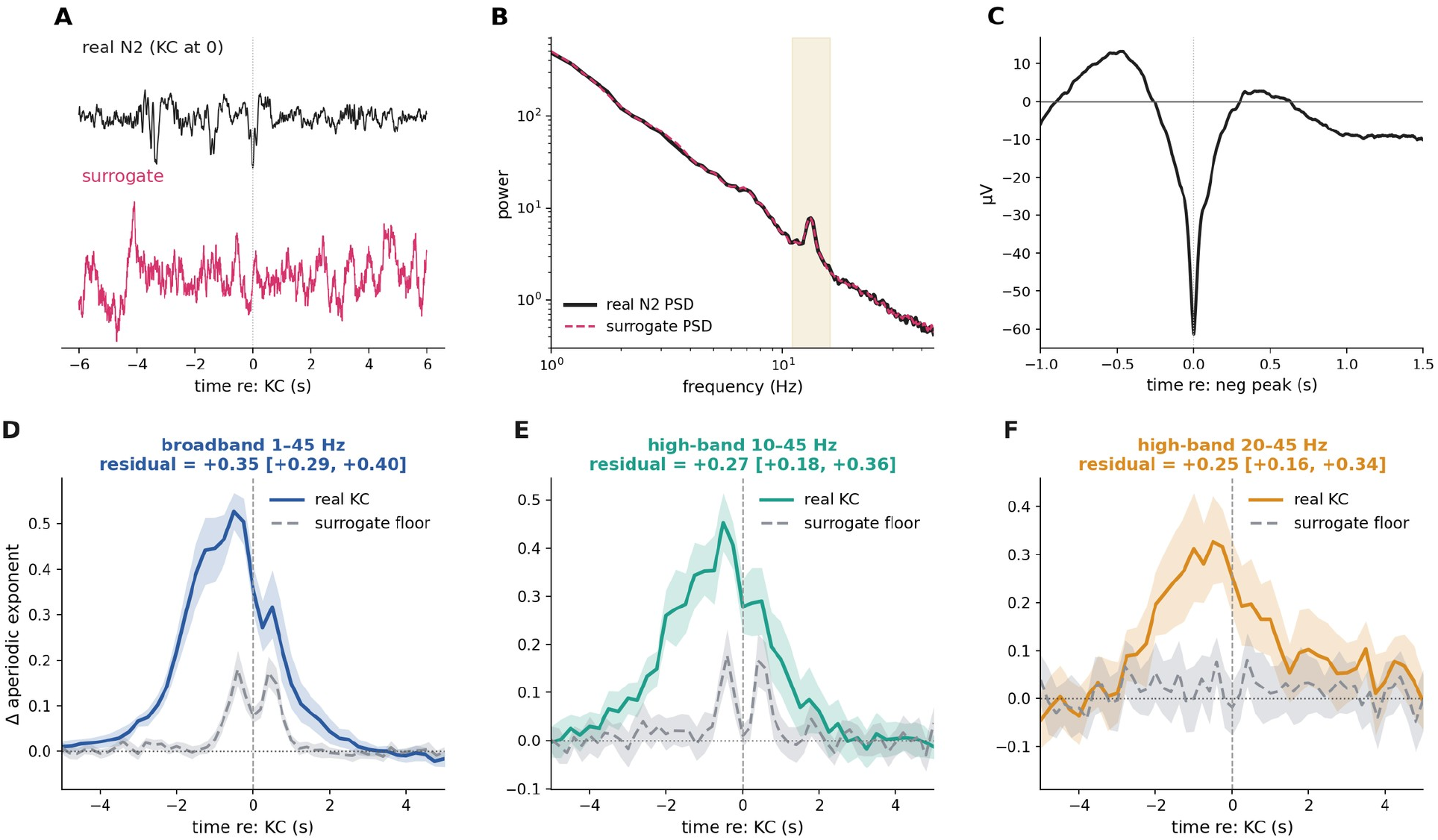
Isolating genuine aperiodic change from waveform contamination. (A) Representative endogenous N2 segment containing a KC (top, black) versus a phase-randomized surrogate of the same segment (bottom, pink). (B) PSD of the real N2 signal (black) versus its surrogate (dashed pink), confirming the surrogate preserves the real signal’s spectral content (sigma/spindle band, 11-16 Hz, shaded). (C) Empirical KC template, built by averaging isolated, expert-annotated KC waveforms aligned to the negative peak and baseline corrected, used to inject artificial events into the surrogate. (D-F) Event-locked changes in aperiodic exponent dynamics for real KCs (solid) versus the injected template surrogate floor (dashed gray line) in the broadband (D) and highband, 10-45 Hz (E) and 20-45 Hz (F), fit ranges. In all three ranges, the real deflection exceeds the surrogate floor, leaving a residual not attributable to the waveform (broadband Δχ ≈ 0.35, 95% CI [0.29, 0.40]; 10–45 Hz Δχ ≈ 0.27, [0.18, 0.36]; 20–45 Hz Δχ ≈ 0.25, [0.16, 0.34]; all p < 0.001).

To test how waveform variability influences aperiodic dynamics, each KC’s negative-peak amplitude was correlated across events with its exponent deflection (Figure 4A-C). In the broadband fit, larger KCs produced larger deflections (mean within-subject r = 0.39 ± 0.11, across subjects t(18) = 16.0, p<0.001; Figure 4A), suggesting waveform contamination and biased model fits. This amplitude dependence was strongly reduced in the 10-45 Hz highband fit (r = 0.10 ± 0.12, t(18) = 3.6, p = 0.002; Figure 4B) and absent in the 20-45 Hz highband fit (r = 0.02 ± 0.09, t(18) = 1.1, p = 0.27; Figure 4C). The attenuation of this relationship across the highband fits (Figure 4D) indicates that the amplitude-driven effect is largely confined to low frequencies. Tracking coupling between negative-peak amplitude and exponent deflections as a function of the lower fit bound showed a steep decay between ∼2 and 6 Hz (Figure 4E). A residual relationship remained at 10-45 Hz (r = 0.10 ± 0.11, t(18) = 3.6, p = 0.002). Nonetheless, the strong attenuation of the amplitude coupling in the highband is consistent with the surviving component reflecting a genuine change in the aperiodic background.

**FIGURE 4.**
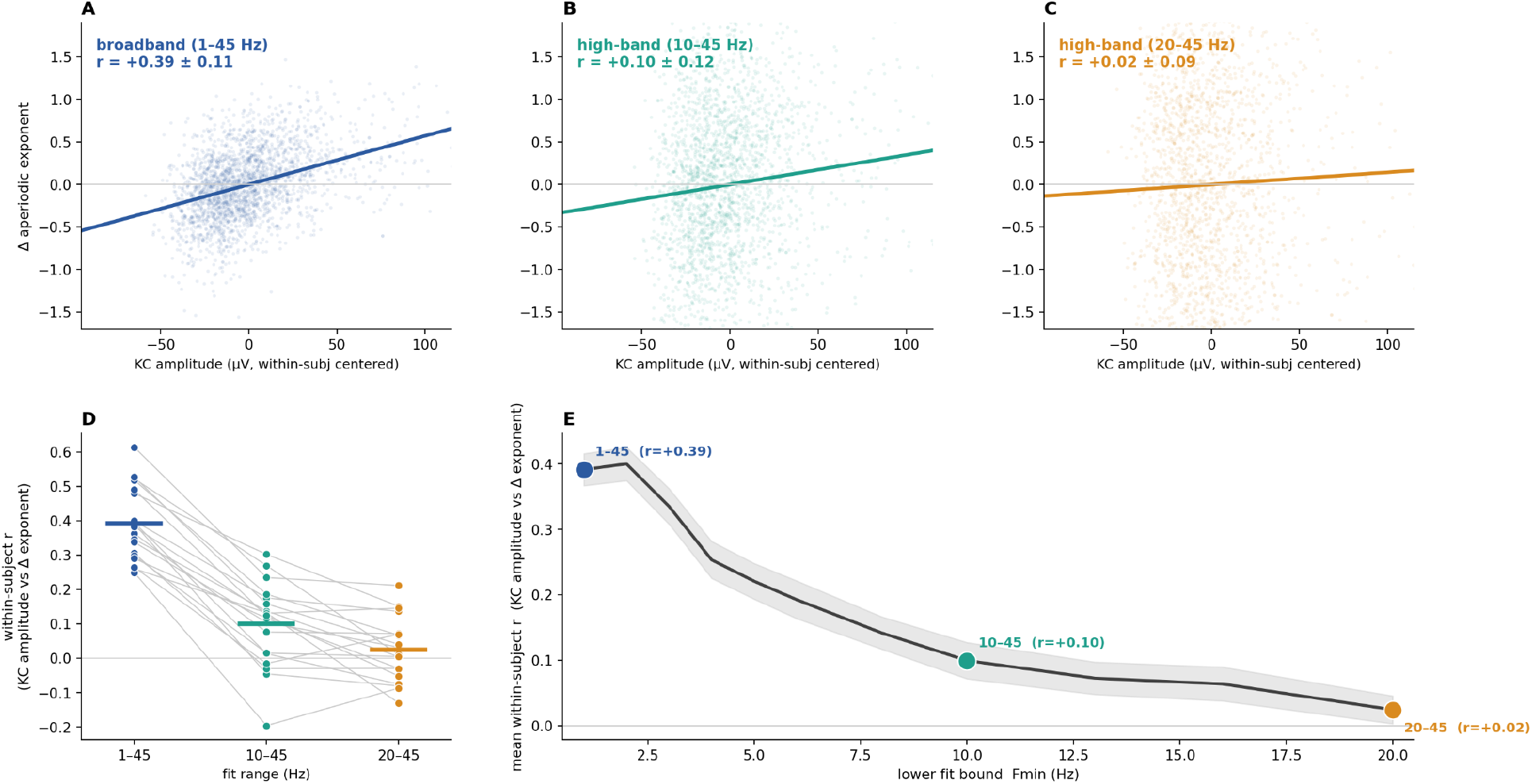
Waveform amplitude dependence of the exponent deflection across fit ranges. (A-C) Δχ plotted against KC negative-peak amplitude for individual events, pooled across subjects, for the broadband (A) and highband (B&C) fits. Amplitudes of each event were within-subject centered, relative to that subject’s mean, to remove between-subject differences in KC size. The positive relationship in the broadband fit (mean within-subject r = +0.39 ± 0.11, t(18) = 16.0, p < 0.001) is progressively attenuated at 10–45 Hz (r = +0.10 ± 0.12, t(18) = 3.6, p = 0.002) and absent at 20–45 Hz (r = +0.02 ± 0.09, t(18) = 1.1, p = 0.27). (D) Within-subject amplitude versus deflection correlation across three fit ranges. Individual subjects are connected by gray lines, showing consistent within-subject decline. (E) Mean within-subject correlation as a function of the lower fit bound (Fmin), showing a steep decay between ∼2-6 Hz before plateauing near zero, indicating the amplitude-dependent component is concentrated at low frequencies.

As a final control, we removed the KC waveform from each window directly and re-estimated the aperiodic exponent dynamics (Gyurkovics et al., 2022). Regressing the empirical KC template left the highband deflections unchanged (Figure 5; 10–45 Hz: Δχ = 0.43 before versus 0.41 after regression; 20–45 Hz: Δχ = 0.30 versus 0.29). In each band the after-regression deflection remained significantly above zero (10-45 Hz: t(18) = 16.1; 20-45 Hz: t(18) = 12.7, both p < 0.001), indicating that these increases are not dependent on the waveform. Contrastively, the broadband deflection reduced modestly (1–45 Hz: Δχ = 0.51 before versus 0.43 after regression), though it remained significantly above zero (t(18) = 21.4, p < 0.001). That all three residual deflections stayed significantly positive suggests the exponent increase reflects a change in the aperiodic background rather than the waveform shape. These findings converge with the fit-range, amplitude, and surrogate-floor controls.

**FIGURE 5.**
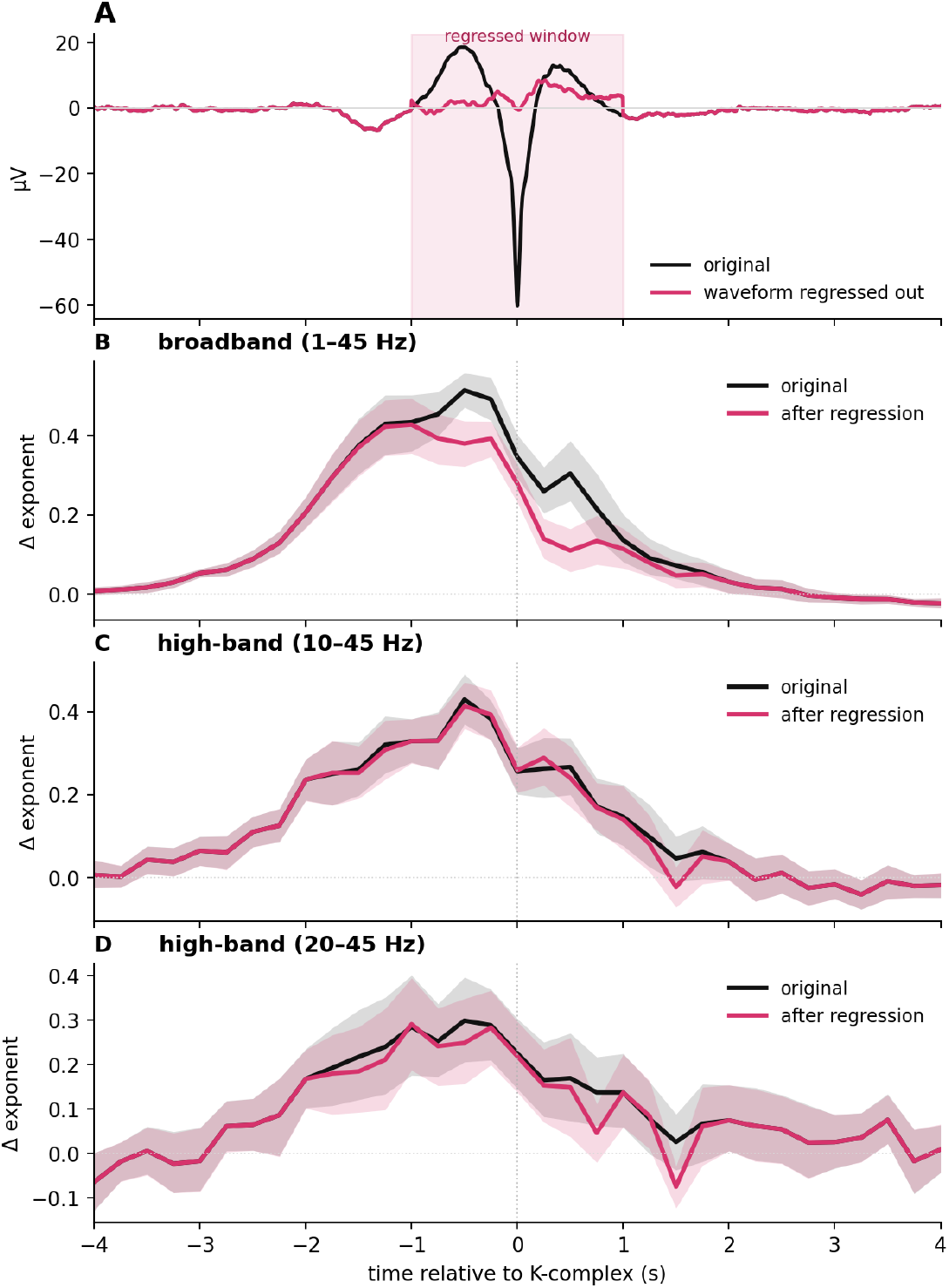
Exponent increases in high-band fits survive KC waveform regression. (A) Grand-average voltage around KC, before (black) and after (pink) regressing the empirical KC template out of the +- 1 second window at the negative peak (shaded); the deflection is removed inside the shaded window while the surrounding activity is unchanged. (B-D) Event-locked aperiodic exponent, relative to N2 baseline, computed on the original signal (black) and after the waveform regression (pink), for the broadband (1-45 Hz) and highband (10-45 and 20-45 Hz) fits. Removing the waveform reduces the broadband deflection (B) but leaves the highband deflections essentially unchanged (C,D), indicating that the highband increase reflects a change in the aperiodic background rather than distortions introduced by the waveform.

## 4 DISCUSSION

Aperiodic neural activity has been shown to reflect physiological processes that are dynamically modulated across and within vigilance states (Lendner et al., 2020; Ameen et al., 2024). We hypothesized that the KC, given its proposed roles in sleep stage regulation, cortical down-state production, and memory consolidation, is accompanied by distinct aperiodic dynamics (Halász, 2005; Cash et al., 2009; Helfrich et al., 2021). Quantifying these dynamics around the sharp, large amplitude deflections of the KC, especially at sufficient temporal resolution, introduces artifacts in the frequency-domain that must be carefully considered (Cellier, 2026).

The present study examined whether KCs are accompanied by a genuine change in aperiodic activity, or whether apparent changes instead reflect the KC waveform distorting model fits of the 1/f exponent. Using time-resolved spectral parameterization of overnight sleep EEG, we first characterized the aperiodic dynamics locked to expert-annotated KCs. To disentangle cortical state dynamics, reflected in the exponent of the power spectrum, from spectral artifacts introduced by the KC waveform morphology, we applied three novel complementary controls: a spectral fit restricted above the KC’s frequency footprint, an injected-waveform artifact floor, and an amplitude-modulation test across events. We also applied an established method for disentangling true aperiodic differences from distortions introduced by event-related-potential waveforms (Gyurkovics et al., 2022).

Across these controls, a component of the KC-locked aperiodic dynamics was independent from the waveform’s contribution. The positive exponent deflection was minimally reduced when using high-band frequency ranges to fit the model to the spectrum. Across all fit ranges, the deflection in real data was significantly greater than that of simulated events derived from combining surrogate background activity with template waveforms. The deflection was weakly correlated with KC amplitude when using broadband fits, and uncorrelated when the model was fit at higher frequency ranges. Finally, directly regressing the KC waveform out of each window left the high-band deflection unchanged and only modestly reduced the broadband deflection, with all residuals remaining significant. Because this approach removes the waveform directly rather than restricting the fit range or simulating a background, it provides converging evidence that does not depend on the assumptions of the novel controls. Together, these results indicate that KCs are accompanied by a genuine steepening of the aperiodic 1/f exponent.

These findings are consistent with the current understanding of the physiology and function of KCs established in previous literature. Both evoked and spontaneous KCs recorded at the scalp are accompanied by an influx of hyperpolarizing current and decreased neuronal firing characteristic of a cortical down-state (Cash et al., 2009), which could function to suppress sensory information and protect sleep (Halász, 2005). The steepening of the aperiodic exponent during KCs may reflect a non-invasive index of this state shift (Gao et al., 2017) or, more generally, an increase in population spiking synchrony (Preston et al., 2026). These findings align with the role of KCs in the gating of sensory processing and may implicate KCs in the modulation of thalamocortical activity and therefore network communication and memory consolidation (Mak-McCully et al., 2015), although additional investigation into aperiodic activity’s influence on coupled sleep oscillations is needed.

These results also highlight the challenges of separating discrete events from background activity, especially when the two are correlated. This was recently demonstrated through the observation of auditory-evoked aperiodic dynamics independent of a KC waveform (Ameen et al., 2025), as well as the automated detection of putative sharp-wave-ripples in purely aperiodic signals (van Schalkwijk and Helfrich, 2026). However, because common detection algorithms use adaptive thresholds for event detection, an aperiodic-only surrogate lowers the detection threshold and inflates false positives, whereas recent work showed that a spectrum- and amplitude-matched surrogate does not (Kragel, 2026). These opposing views share a common limitation. Both are argued from aggregate detection rates across conditions rather than from individual events, so neither can establish whether a particular event is genuine or aperiodic. In the present study, we instead address the question at the level of individual events. Rather than asking whether events are detected in a surrogate, we ask how much of a real, individually identified event’s spectral changes the waveform can produce on its own. Our surrogate floor control applies the same standard Kragel’s critique demands: that a fair null must reproduce the real signal’s structure, inx this case its full power spectrum, rather than aperiodic activity alone. Our approach allows us to address the question that the detection-rate debate leaves open, whether a given event reflects genuine dynamics or the background against it is measured.

More broadly, our approach offers a strategy for separating discrete events from the correlated aperiodic activity they may arise within, wherever the two are difficult to disentangle.

Further, these results emphasize the importance of selecting model parameters in time-resolved spectral analysis. Ameen et al. (2025) demonstrated that the choice of frequency range and aperiodic mode significantly alters the derived exponent, and Cellier (2026) explained how the selection of time window duration, oscillatory peak count, and across-trial averaging techniques can influence the resolution and accuracy of model fits. The present results reiterate this message by demonstrating how aperiodic estimates can be distorted by discrete waveforms, and that this distortion itself is modulated by model parameters. This parameter-dependence underscores that model parameters should be constrained primarily constrained by the empirical question that the model is being used to answer, including tuning the time-frequency tradeoff to the resolution of the dynamics of interest and using aperiodic and periodic settings that most accurately represent the underlying physiology.

The present findings contribute to a growing body of work suggesting that aperiodic activity carries meaningful physiological information and plays a functional role in brain networks (Gao et al., 2017; Pei et al., 2023; Lu et al., 2024; Monchy et al., 2025; Preston et al., 2026; van Engen et al., 2026). Aperiodic dynamics appear to index ongoing fluctuations in cortical state that contribute to sleep macro- and microarchitecture. This interpretation may provide insight into previous findings of altered KCs across neurological and psychiatric disorders. Many of these conditions have independently been associated with disruptions in E/I balance and large-scale cortical dynamics (Žiburkus et al., 2013; Gonçalves et al., 2017; Sohal and Rubenstein, 2019; Lányi et al., 2024; Kovacevic et al., 2025; Li et al., 2025). Consequently, abnormal KC activity may not reflect abnormalities in isolated event generation, but shifts in underlying neural state dynamics, consistent with existing work characterizing changes in E/I balance during sleep (Chellappa et al., 2016; Niethard et al., 2016; Brécier et al., 2022; Brodt et al., 2023).

Several limitations should be noted. The presented dataset contained subjects within a narrow age range and therefore precluded investigation of age-related changes, known to impact both KCs and aperiodic activity (Crowley et al., 2002; Hill et al., 2022; McSweeney et al., 2023; Leroy et al., 2025). Further, event detection in these analyses was restricted by expert annotation.

While often treated as the gold standard, expert annotation is outdated, subjective, and unreliable, and future analyses should rely on improvements in data-driven detection methods (Lajnef et al., 2017). Additionally, cortical oscillations during sleep can be globally synchronized, occur locally, or propagate as traveling waves (Massimini et al., 2004; Nir et al., 2011; Mak-McCully et al., 2015). Our single-channel analyses are unable to resolve these distinct spatiotemporal patterns, which may be accompanied by distinct aperiodic dynamics. Lastly, the artifact-floor and template controls depend on the empirical KC template and on the alignment to the negative peak, which deliberately maximizes waveform contamination. Therefore, the surviving component should be interpreted as a conservative estimate of the genuine aperiodic change.

In summary, we demonstrate that KCs are accompanied by a phasic increase in the aperiodic exponent that cannot be completely attributed to spectral contamination by the event waveform. Our findings suggest that the KC and sleep microarchitecture in general cannot be fully characterized by discrete oscillatory events alone. Rather, it is likely that the interactions of these events with the underlying aperiodic dynamics are necessary for supporting the diverse network-level functions of sleep. While it is tempting to say – based on our observation that realistic looking KCs can arise from purely aperiodic dynamics (Figure 1) – that KCs are artifacts of seeing structure in noise, it is clear that aperiodic dynamics do give rise to reliable KC events. However, our approach highlights a methodological challenge in disentangling physiological changes from artifacts introduced by transient waveforms in time-resolved spectral analyses, and we introduce a set of complementary controls as a potential solution. To extract the most accurate representations of the underlying physiology, spectral parameterization techniques must carefully consider all sources of power introduced by the multitude of simultaneously occurring neural events. At the same time, we must acknowledge that not every discrete event – be it an oscillatory burst, a KC, or a sleep spindle, even when expertly annotated – necessarily represents anything more than aperiodic dynamics. Thus, as time-resolved spectral parameterization becomes routine, distinguishing genuine shifts in aperiodic activity from the spectral signatures of transient waveforms will be essential to correctly interpreting these dynamics.

## Acknowledgements

We would like to acknowledge McKena Geiger, Mhone Bogonko, and Jonathan Ahern for their feedback on the manuscript. This work was supported by the National Institute of Mental Health grant R61MH135109 to B.V.

## 5 SUPPLEMENT

**Supplementary Figure 1:**
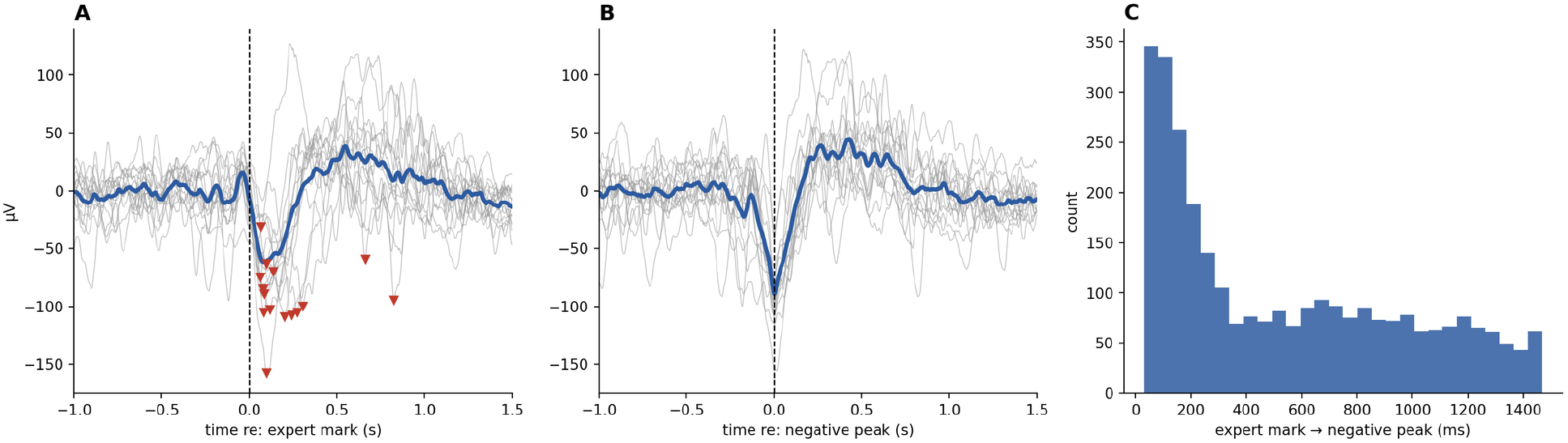
Variability in expert annotations from MASS dataset. (A) 15 KCs from one randomly selected subject aligned to the expert mark (t=0; individual events, grey; mean, blue; red triangles indicate each event’s negative peak. (B) The same events aligned to the negative peak. The averaged waveform produces a sharper negative peak, more aligned with defined KC morphology. (C) Distribution of the time from the expert annotation to the negative peak across 2,937 KCs (six randomly selected subjects); there is large variability in the annotation position (SD 430 ms, IQR 142-879 ms).

